# WxS-QC - a quality control pipeline for human Whole-Genome and Whole Exome sequencing cohorts

**DOI:** 10.64898/2025.12.15.694327

**Authors:** Gennadii Zakharov, Ruth Y. Eberhardt, Alina Frolova, Dmytro Horyslavets, Eric Hidari, Iaroslav Popov, Joanna Dennehy, Mykhailo Koreshkov, Pavlos Antoniou, Valeriia Kantsypa, Vladimir Ovchinnikov, Vivek Iyer

## Abstract

**Motivation:** Whole-exome (WES) and whole-genome (WGS) sequencing are rapidly becoming preferred methods for population-scale analysis of the human genetic landscape. However, there are currently no standardized quality control (QC) pipelines for human WES and WGS datasets. Moveover, there are no open datasets that can be used to test QC pipelines, because most projects (like 1000 genomes and gnomAD) publish only post-QC results.

**Results:** We present WxS-QC, a powerful, scalable, and convenient pipeline for the QC of human WGS and WES cohorts, developed at the Wellcome Sanger Institute (WSI). Our pipeline is based on a deep refactoring of the gnomAD quality control pipeline code and is aligned with current best practices in WGS/WES cohort data QC. It offers a set of novel QC techniques, automatic export of resulting graphs and summary tables, excellent performance and scalability, and incorporates comprehensive documentation. To test our pipeline and similar solutions, we also assembled an open dataset with all required metadata.

**Availability and implementation:** The pipeline code is written in Python using the Hail library and is freely available under the BSD-3 license here: https://github.com/wtsi-hgi/wxs-qc. It can run in any UNIX-like environment and has been able to efficiently process cohorts of up to 200,000 whole-exome samples with the potential to handle bigger datasets, depending on the available hardware. The detailed description of the pipeline, resources and test data is available in the pipeline documentation: https://github.com/wtsi-hgi/wxs-qc/blob/main/README.md

The open test dataset is available to download from https://wxs-qc-data.cog.sanger.ac.uk/wxs-qc_public_dataset_v3.tar. An example of test dataset analysis is available in the supplementary materials.

## Introduction

The advent of high-throughput sequencing technologies has revolutionised biological research and clinical diagnostics. Whole-exome sequencing (WES) and whole-genome sequencing (WGS) are rapidly becoming preferred methods for population-scale association studies of complex traits (Sealock et al. 2025). Efforts focusing on collecting and harmonizing sequencing data range from large cohorts such as the Genome Aggregation Database (Chen et al. 2024) and UK BioBank (Van Hout et al. 2020) to smaller disease-specific or ethnically specialised cohorts (Deciphering Developmental Disorders Study 2015; Sazonovs *et al*. 2022; Koko *et al*. 2024; Kim *et al*. 2025).

Prior to being subjected to statistical analysis, every cohort requires a careful quality control process to remove poor-quality samples and to distinguish true variants from technical artifacts (Sealock et al. 2025). Despite the rapid development of software tools for genome data analysis, there is currently no standardized WES and WGS data quality filtering pipeline for the purpose of population-scale association analysis (Sealock *et al*. 2025).

One of the recent attempts to fill this gap is the tutorial (Sealock *et al*. 2025), which provides guidelines for conducting QC of samples and variants, and compares commonly used software programs for quality control. The tutorial mentions PLINK (Purcell *et al*. 2007), VCFtools (Danecek *et al*. 2011), and BCFtools (Danecek *et al*. 2021) as widely used utilities for performing WES and WGS data QC. However, these tools cover only some specific aspects of data QC, and require manual work to achieve the goal. The tutorial authors suggest using the Hail library (Poterba *et al*. 2025) due to its scalability and flexibility, to provide a simple pipeline for genome/exome data QC. However, this simple pipeline uses hardcoded paths and constants for configuration, and provides no visualisations or export of results. Furthermore, it does not have an independent variant QC module and relies on external GATK VQSR scores (Auwera and O’Connor 2020) to perform variant QC.

A much more powerful and comprehensive solution is provided by the codebase used for data QC of the gnomAD database (Chen *et al*. 2024). This code generally aligns with the tutorial (Sealock *et al*. 2025) for sample QC and uses the Random Forest algorithm to perform variant QC. However, this codebase is tightly coupled to the Broad Institute infrastructure and data, which makes it challenging to use outside the Broad Institute. The gnomAD codebase also contains minimal reporting and visualization functionality.

As a result, there is currently no open solution for a powerful, scalable, and convenient human genome/exome cohort QC.

## Results and Discussion

In this paper, we present **WxS-QC**, a pipeline for WGS/WES cohort QC. Key advantages of this pipeline are:

1. It is based on a deeply refactored version of the gnomAD v3 quality control pipeline (Chen *et al*. 2024) with some backports from gnomAD v4, and aligned with best practices described in (Sealock *et al*. 2025). We based our pipeline on the gnomAD v3 codebase because gnomAD v4 has been augmented to work with very large-scale datasets (several million exomes), and so implements techniques that are not yet required for the majority of WES/WGS cohorts, which have a more moderate scale. We plan to adapt gnomAD v4 in the near future to enable processing of cohorts with up to several million exomes.
2. It is easily configurable via a YAML-based file and contains no hardcoded paths or constants.
3. It incorporates novel QC techniques validated by WSI and external scientific teams working with human WES and WGS cohorts (Koko *et al*. 2024; Kim *et al*. 2025). In particular:
  a. It uses a PCA projection approach to make superpopulation prediction robust (see below).
  b. It adds a “genotype hardfilter evaluation” step, which allows the user to estimate the effect of different filter threshold combinations. Users can choose an optimal combination of variant and genotype-level filters to suit their scientific goal.
4. We provide a comprehensive, user-friendly tutorial describing the analysis process along with an open test dataset. The complete example of the QC for the public test data we provided is described in the supplementary materials.
5. The pipeline automatically exports summary tables and interactive graphs, so the user does not have to figure out which summaries are useful or how to plot them.
6. The pipeline uses Hail (https://github.com/hail-is/hail) – an open-source library for large-scale genomic analysis designed to handle massive datasets (Poterba *et al*. 2025). The pipeline has been tested on cohorts of 170,000 whole-exome samples and 770 whole-genome samples.
7. The pipeline is infrastructure-agnostic and can run in any environment that supports Hail and Spark: a local machine, an on-premises Spark cluster, or a commercial cloud.

### Pipeline scheme

The high-level pipeline schema is shown in Figure 1.

1. The pipeline expects input VCFs called by GATK v4 (Auwera and O’Connor 2020): we suggest taking VCFs before applying GATK VQSR because our pipeline uses a Random Forest-based method to perform variant QC. The pipeline can also accept optional metadata: contamination check provided by VerifyBamID2 (Zhang *et al*. 2020), self-reported sex, pedigree information, etc. The complete set of metadata is described in the pipeline documentation: https://github.com/wtsi-hgi/wxs-qc/blob/main/docs/wxs-qc_howto.md
2. The **metadata validation step** converts raw VCFs to Hail data structures and validates them against the provided metadata.
3. The **sample QC** step broadly follows the approach described in (Sealock *et al*. 2025): we subset each sample for high-quality variants, impute the sex for each sample (and check consistency with reported sex), conduct a principal component analysis (PCA) to correctly assign a superpopulation for each sample (Patterson, Price and Reich 2006; Reich, Price and Patterson 2008), and apply the superpopulation-stratified threshold to filter samples. To predict super populations, our pipeline uses the PCA projection approach - namely, we run PCA on samples drawn from the 1000 genomes study (1000 Genomes Project Consortium *et al*. 2015) and then use Hail’s `pc_project()` function to assign superpopulations to the samples from the study. This approach ensures that PC axes remain stable and not distorted by unusual or specific ancestry, because they are always defined by the same reference set of samples panel (Wang *et al*. 2015; Zhang, Dey and Lee 2020). As a result, our pipeline is independent of the distribution of samples amongst the superpopulations and can handle datasets with any number of related individuals without needing to remove related samples, and the PCs from different studies are directly comparable.
4. The **variant QC step** uses the approach used for gnomAD v2 (Karczewski *et al*. 2020) and the first stages of gnomAD v3: we use a set of open resources to label variations as likely True-Positives and likely False-Positives, train a random forest (RF) model on these data, and use the RF model score to group variants into several bins based on their reliability.
5. The final module, hard filter evaluation and genotype QC, is a new functionality developed at the Wellcome Sanger Institute. The detailed description of this step is available in the pipeline documentation and in the paper (Koko et al. 2024). In this step, we test how different combinations of variant-level and genotype-level filters affect the total resulting genotypes and variants. The step also provides plots to allow scientists to review the results and choose optimal combinations.

**Figure 1.**
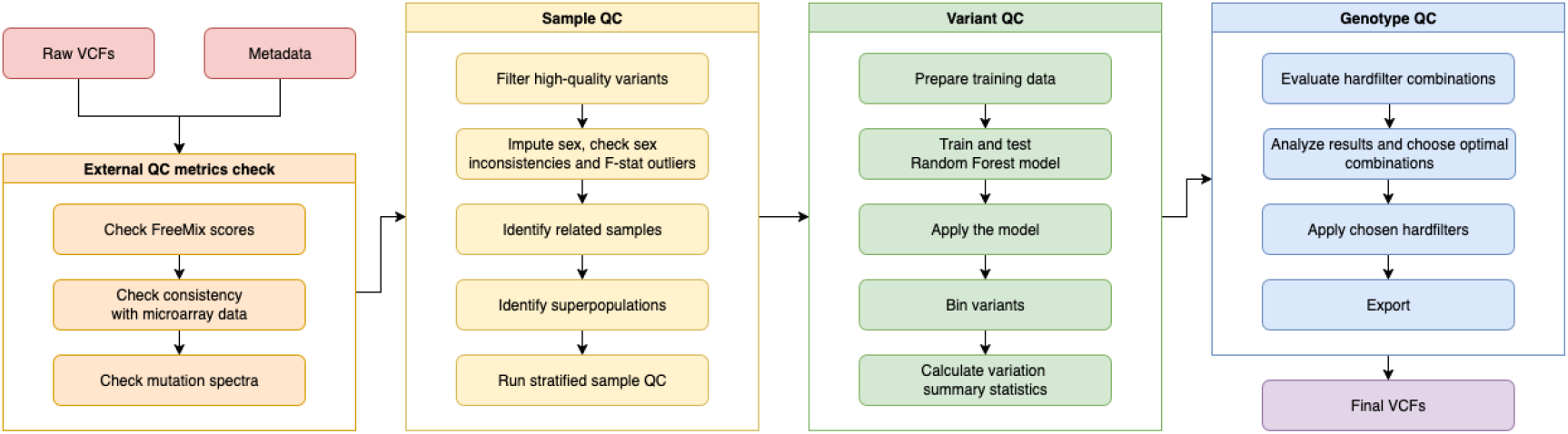
The high-level scheme of the WxS QC pipeline.

## Conclusion

The WxS-QC pipeline, developed by WSI, is a comprehensive and easy-to-configure tool that retains the power of the original gnomAD QC pipeline while adding novel QC approaches, documentation, visualisations, open test data, and infrastructure-agnostic operation.

In the supplementary materials, we demonstrate QC analysis for the example dataset, analyse and discuss the results. We show that the WxS-QC pipeline achieves higher precision levels than standard threshold-based filtering, with comparable recall levels, particularly for indels.

We believe that this pipeline is helpful for anyone who runs sequencing of human genome/exome data.

## Supporting information

Exome cohort QC with the Sanger WxS-QC pipeline

## Code and data availability

- The pipeline code and documentation are freely available via the WSI GitHub repository: https://github.com/wtsi-hgi/wxs-qc
- The resource bundle is available to download from https://wxs-qc-data.cog.sanger.ac.uk/wxs-qc_resources.tar
- Open test dataset is available to download from https://wxs-qc-data.cog.sanger.ac.uk/wxs-qc_public_dataset_v3.tar

## Conflicts of interest

Authors declare no conflicts of interest.

## Acknowledgements

The authors sincerely thank Klaudia Walter, Mahmoud Koko, and Hilary Martin for their invaluable support and guidance through the results analysis and approach improvements that made this publication possible.

## Funding information

This research was funded in whole, or in part, by the:

- Wellcome Trust 220540/Z/20/A. For the purpose of Open Access, the author has applied a CC BY public copyright licence to any Author Accepted Manuscript version arising from this submission.
- MRC Exome Sequencing of ALSPAC, Born in Bradford and Fenland Cohorts (reference MC_PC_23015).
- Research Collaboration Agreement L24-0442/0124U004344 (22-August-2024) between Genome Research Limited (UK) and the Institute of Molecular Biology and Genetics of National Academy of Sciences of Ukraine (IMBG).

## Supplementary data

Exome cohort QC with the Sanger WxS-QC pipeline.

## Notes

https://github.com/wtsi-hgi/wxs-qc

### Competing Interest Statement

The authors have declared no competing interest.

