## Supplementary material for "WxS-QC - a quality control pipeline for human Whole-Genome and Whole Exome sequencing cohorts": Exome cohort QC with the Sanger WxS-QC pipeline

### Supplementary 1. Exome cohort QC with the Sanger WxS-QC pipeline

#### Intro

Here we provide an example of exome cohort QC using the WxS-QC pipeline. For input, we are using the publicly available example dataset that we constructed from the open 1000 genomes data (1000 Genomes Project Consortium *et al.* 2015), published with the pipeline (referred to in the subsequent text as “the example dataset”). All the details regarding the construction of the example dataset are described at the end of this text and in the relevant GitHub repository. This analysis is based on pipeline version 0.8. To use the most recent version of the pipeline, please refer to the upstream version of the pipeline howto.

##### Set up

The Sanger WxS-QC pipeline is cloud-agnostic and can be run on any UNIX-compatible machine running Python 3 and Hail. However, please note that the Hail data structures are “heavy” in terms of storage space and memory. Before starting, ensure you have enough memory (for acceptable speed, at least 0.25x the size of the input VCFs) and storage space (at least 10x the size of the input VCFs) to proceed.

The pipeline setup process is described in the setup documentation. Briefly, you need to do the following:

1. Clone the Sanger WxS-QC repository.
2. Switch to a specific version of the pipeline if you need to compare cohorts between runs. This example was done using version 0.8.
3. Set up the Python virtual environment following the environment setup howto .
4. Download the example dataset as described in the pipeline howto .
5. Create the configuration file for the dataset and set up the analysis folder as described in the pipeline howto. To repeat this analysis, simply change the root path to the analysis folder.
6. Download the resource bundle, unpack it and place it in the analysis folder.

The pipeline is then ready to run.

#### Analysing the data

Detailed instructions on how to run the pipeline and obtain the results are available in the pipeline howto. In this text, we focus on the QC results and interpretation. The pipeline saves all human-readable results in the “`annotations`” folder and puts all graphs in the “`plots`” folder. All of the results below are derived from the example dataset that can be downloaded from https://wxs-qc-data.cog.sanger.ac.uk/wxs-qc_public_dataset_v3.tar as specified in the pipeline howto.

### Stage 0. Preparing resources

First, prepare the resource files needed by following the WxS-QC resources how-to. Then run steps 0.1 and 0.2 in the pipeline how-to to convert resource files into Hail structures. If you work with whole-genome sequences and want to use whole-genome gnomAD frequencies, refer to the resources preparation description.

Once the resource files have been converted to Hail structures, this step doesn't need to be repeated for every dataset. When you run a new analysis, you can symlink the Hail tables to the new matrixtable folder. Check that all steps have been completed without errors and proceed to the next step.

### Stage 1. Data import and annotation

Run all steps from stage 1 to import and annotate data, according to the pipeline howto. At this stage, you need to review the following results.

#### Freemix scores

The FreeMix score, produced by the VerifyBamID (Zhang et al., 2020) utility, represents the level of contamination in the sample. The pipeline reads this file, makes the plot and identifies the problematic samples. Review the graph "`freemix_validation.html`" (Figure 1). All samples from the example dataset pass the FreeMix score threshold <=0.05.

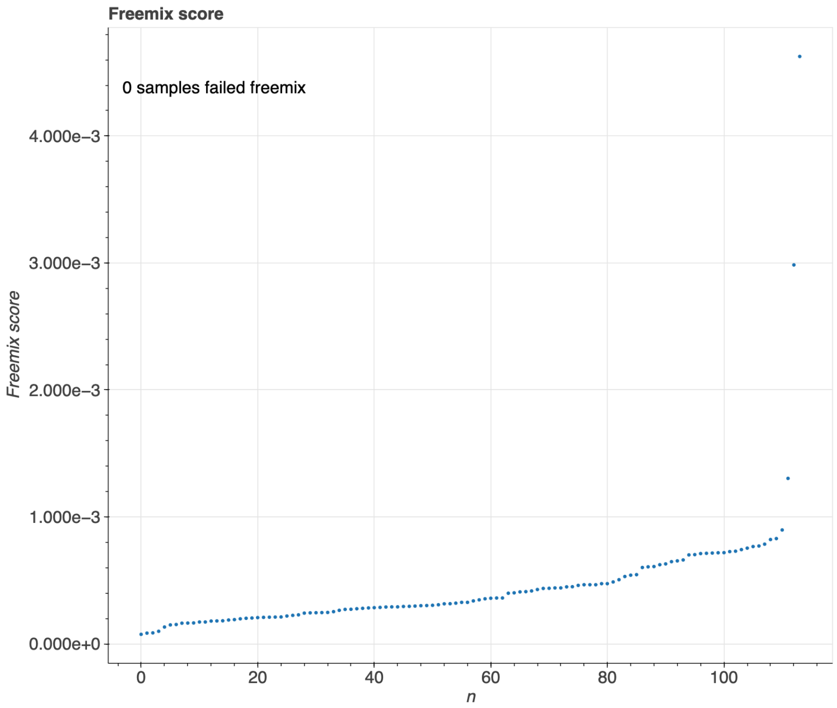


Figure 1. Freemix score validation for the example dataset samples. The X-axis contains the sample number. Samples are sorted by the Freemix score (Y axis)

All the samples failing the Freemix score are reported in the file "`samples_failing_freemix.tsv`". We suggest excluding all these samples. To keep Freemix outliers in the dataset, set the ***remove_freemix_outliers*** config option to "false".

***Note***: If a significant part of the samples in your dataset (>5%) is contaminated, it can affect the sample QC. In this case, we suggest that you manually remove contaminated samples from the input VCF(s) and rerun the pipeline from the data import stage.

#### Mutation spectra

Review the graph "`mutation_spectra_preqc.html`". Your mutation spectra should resemble the spectra of the example dataset (Figure 2). The example dataset contains only 114 samples, so we expect significant dispersion and outliers. For bigger datasets, you should have fewer outliers. A significant number of outlying samples can indicate sequencing artefacts.

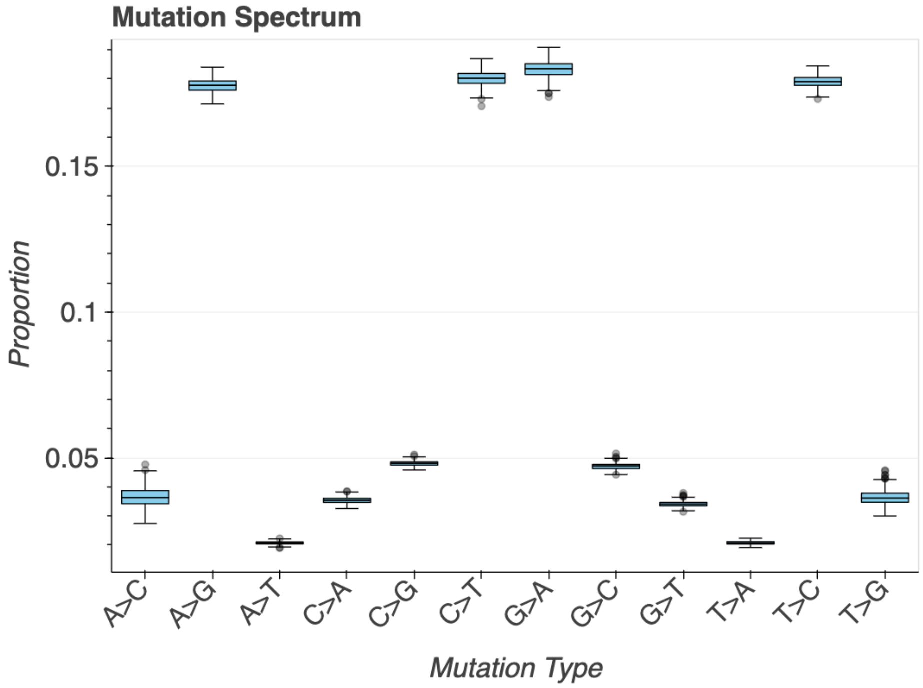


Figure 2. Mutation spectra boxplots for the WxS-QC example dataset before filtering.

If you have a few outlying samples, you can hover your mouse over them to find their IDs and exclude them before the final stage of sample QC.

#### Verification of sample identity

If you have external genotyping data, for example, from microarray genotyping, you can use bcftools gtcheck (Danecek *et al.* 2021) to verify the sample's identity. The WxS-QC pipeline can import results of bcftools gtcheck and flag all potentially mismatched samples. The detailed description of this step and interpretation of the step results are described in the [pipeline howto](pipeline howto).

The example dataset has no microarray validation data, so step 1.3 has no effect.

### Stage 2. Sample QC

Run all steps from the sample QC stage according to the [pipeline howto](). At this stage, you should review the following results.

#### Sex imputation and validation

Step 2.1 imputes genetic sex for all samples and compares it with the self-reported sex, if it was provided at stage 1. The pipeline generates the F-stat histogram "fstat_hist.html" (Figure 3) and reports the number of outliers – samples with an F-stat value between the lower and upper thresholds. Both thresholds are defined in the config file section **step2 -> f_stat_outliers** and are shown on the histogram.

The pipeline saves the list of F-stat outliers (`sex_annotation_f_stat_outliers.tsv`) and samples with inconsistent sex annotation (`conflicting_sex.tsv`). Neither F-stat outliers nor mislabelled samples affect the subsequent QC process. Therefore, in most cases, we keep these samples in the dataset and let the end users decide whether to analyse them or not.

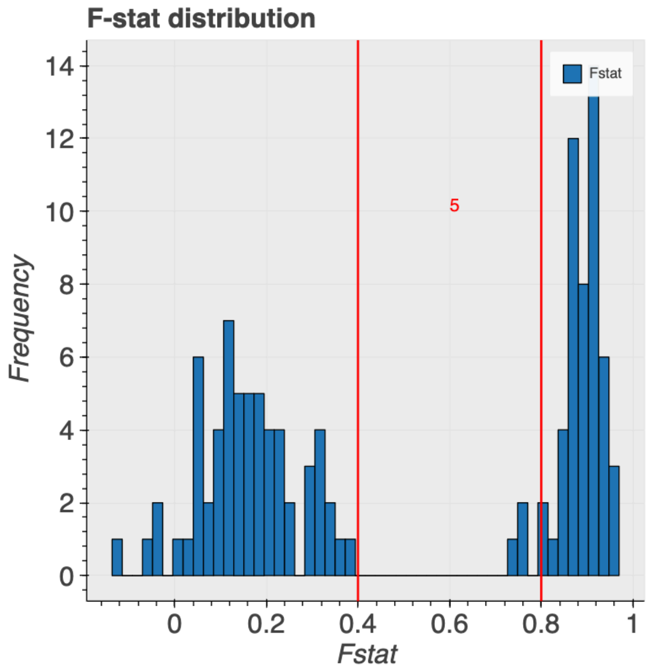


Figure 3. F-stat distribution of the example dataset samples. The low and high F-stat thresholds and the number of outlier samples (between these thresholds) are marked in red.

#### Identify related samples

Step 2.2 performs samples PCA and produces a table of closely related samples (file `annotations/relatedness.tsv.gz`). If you have pedigree information for your dataset, you can check that the step results agree with the pedigree data. The step results don't affect any other pipeline steps.

#### Predict super-populations

Step 2.3 utilizes the PCA projection approach to identify superpopulations. Briefly, we:

- Intersect variation between the example dataset and the set of open variation data from 1000 genomes (1000 Genomes Project Consortium *et al.* 2015). For this study, we used 3202 samples with known superpopulations. The complete list of samples is available together in the dataset archive.
- Run PCA for 1000 genomes samples.
- Project PCA scores onto the example dataset samples.
- Predict superpopulation for the example dataset samples based on projected scores

Since for the PCA we always use 1000 genomes data, our approach works on data cohorts with any superpopulation distribution and incidence of close relatives.

First, review the superpopulation PCA for the 1000 genomes samples (the file `pop_pca_scores_1kg.html`). Figure 4 illustrates the expected appearance of the standard 1000 Genomes PCA graphs.

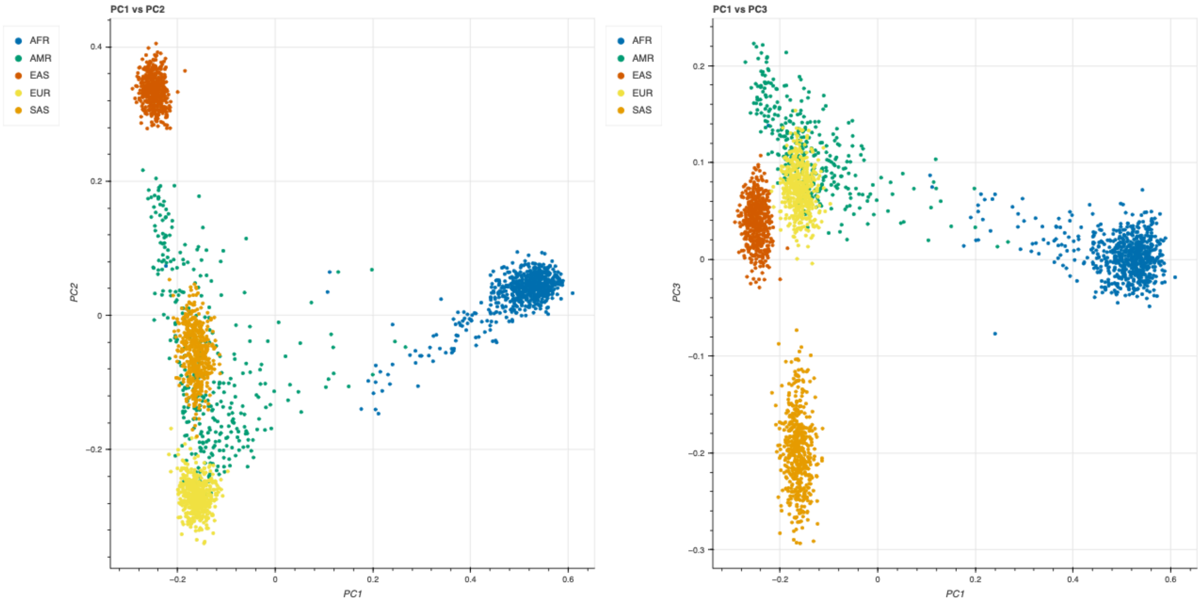


Figure 4. Superpopulation PCA plot for 1000 genomes data.

If everything is Ok, review the PCA results for the dataset (`pop_pca_assigned_scores_dataset.html`). Normally, the dataset samples should be grouped in the same clusters as the 1000 Genomes samples. The number of samples assigned to each superpopulation is available in the console output.

If the superpopulation distribution looks similar to the distribution in Figure 5 (i.e. your superpopulation clusters are co-located with the expected positions of the reference clusters), proceed to the next step.

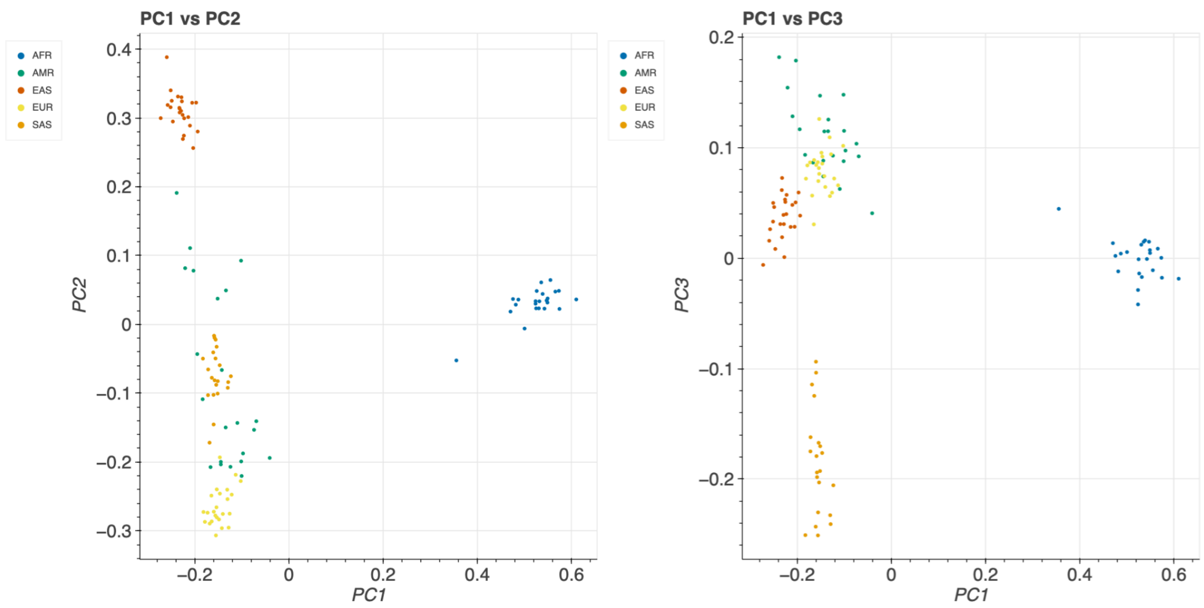

Figure 5. Superpopulation PCA plot for the example dataset.

#### Identify and remove outliers

Step 2.4 runs stratified sample QC using the compute_stratified_metrics_filter() function, identifies outliers and produces sample metric graphs stratified by superpopulation. The `sample_qc_metrics` subfolder in the plots folder contains the file `sample_qc_all_metrics_by_pop.html` with the combined graphs for all metrics and populations. All other files contain larger versions of the same graphs as separate files and are useful for detailed investigation, papers, and reports.

A part of this combined plot for the example dataset is shown in Figure 6. The example dataset contains only 114 samples, so the distribution histograms for each metric look grainy. For big datasets, you should see smoother distributions.

Check the histograms and review that the thresholds and number of outliers are reasonable. Usually, doing this step with the default threshold value of 4 median absolute deviation (MAD) should mark about 1-5% of samples within each superpopulation as outliers. For particular metrics (typically when the superpopulation contains several subpopulations or a small number of samples), you should increase the threshold. The information on how to do this is available in the pipeline howto.

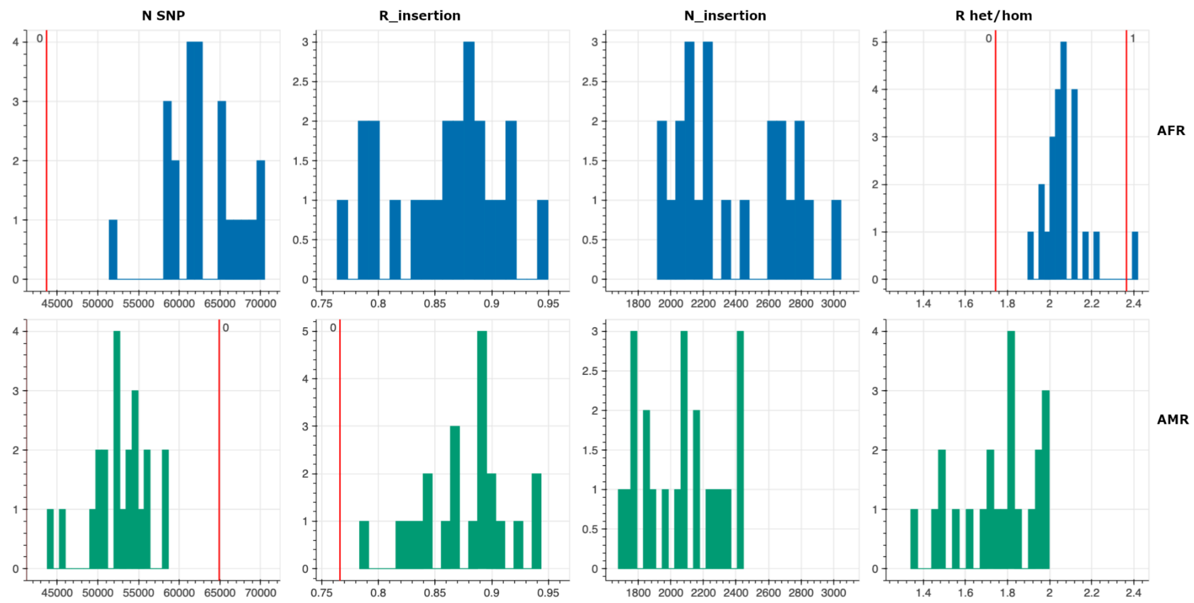


Figure 6. A fragment of the combined SampleQC metrics plot stratified by superpopulation for the example dataset. The corresponding file `sample_qc_all_metrics_by_pop.html` is generated by step 2.4 (Identify outliers) of the WxS-QC pipeline.

##### Compose the final list of samples to remove

Step 2.5 automatically excludes all samples failing at least one metric threshold. If you want to change this behaviour and include outlier samples, save the produced list of outlier samples, set a larger MAD value - to keep all samples inside filtering thresholds - and rerun the outlier identification and remove outliers steps (2.4 and 2.5). Also, for step 2.5, you can provide an additional list of samples you want to exclude from the dataset. If you wish, you can also exclude F-stat outliers or samples with inconsistent self-reported sex. Don't forget to rerun step 2.5 after modifying the list of excluded samples.

#### Stage 3. Variant QC

##### Prepare data for training

In this stage, we prepare a set of variants to train a Random Forest (RF) model predicting variant quality. To do this, we compare our example dataset with known high-quality datasets from the GATK resource bundle (GATK Resource Bundle 2025): 1000 Genomes (1000 Genomes Project Consortium et al. 2015), HapMap (Frazer et al. 2007), Mills and 1000G gold standard indels, and OMNI 2.5 genotypes for 1KG samples and sites. Variants in common between the example dataset and the high-confidence datasets are marked as likely true-positive variants (TP). We also isolate variants in the example dataset with low-quality GATK metrics: QualityByDepth (QD) <= 2, strand bias estimated with Fisher's exact test (FS) >= 60, and RMSMappingQuality (MQ) <=30, and mark them as likely false-positive variants (FP). For details, please refer to the pipeline howto. The same approach is used in gnomAD v2 (Karczewski *et al.* 2020) and gnomAD v3 (Chen *et al.* 2024).

Run the variant QC steps 3.1 and 3.2 according to the [pipeline howto](). Step 3.2 plots histograms of metrics used to mark variants as FP. The graphs for the example dataset are shown in Figure 7.

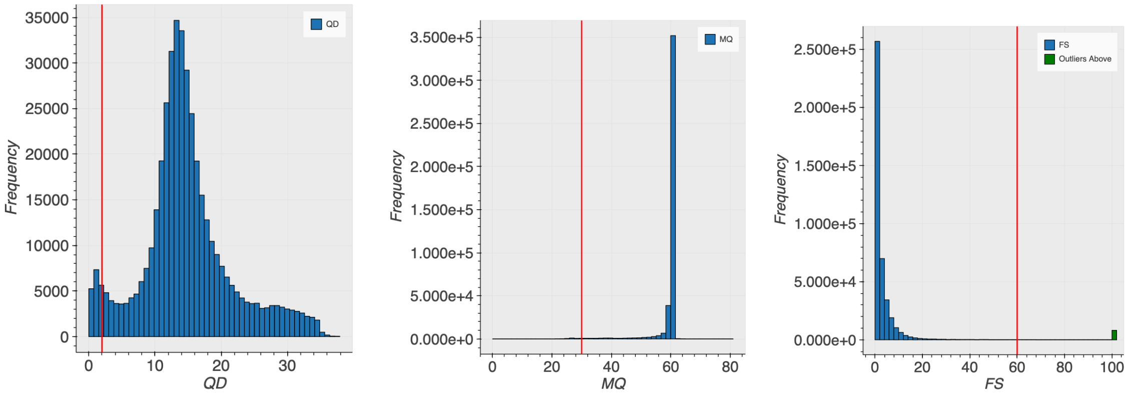


Figure 7. Variant QC metrics used by the QxS-QC pipeline to choose likely false-positive variations. Metric histograms for the example dataset.

Using the default thresholds in the config file we marked 233,357 variations as TP and 23,634 variations as FP in our example dataset. Since TP and FP labels are assigned independently, we have unbalanced TP/FP categories, and we can even have a small number of variants, marked as TP and FP at the same time. The gnomAD random forest function automatically deals with conflicting annotations and balances classes.

#### Train the random forest model

Step 3.3 trains the random forest model and returns a random model ID name. For the example dataset, we use a custom name for convenience, even though this option overwrites the existing model on each re-run. For your own experimental data, we suggest omitting this option and using a random ID. The random ID is generated each time a new model is trained, which improves traceability and allows you to compare and apply different models.

Step 3.3 prints to the console the model ID, overall model accuracy, and the feature importance (Figure 8). All these data are also saved in the file `variant_qc_random_forest/rf_runs.json` under your main analysis folder.

```
{
    "features_importance": {
        "MQ": 0.21330268125259055,
        "MQRankSum": 0.021469220562378787,
        "QD": 0.4198752833512252,
        "ReadPosRankSum": 0.008493531054335566,
        "SOR": 0.07997929763019253,
        "allele_type": 0.0015046316470275 73,
        "has_star": 0.0028365523097713665,
        "meanHetAB": 0.24138679081670011,
        "n_alt_alleles": 0.000900166135977061,
        "variant_type": 0.001972173716637874,
        "was_mixed": 1.0130991694278172e-05,
        "was_split": 0.008269540531468982
    },
    "input_args": {
        "adj": true,
        "transmitted_singletons": true,
        "vqsr_training": false
    },
    "test_accuracy": 0.993519882179676,
    "test_intervals": "chr20"
}
```


Figure 8. Fragment of the model description JSON file, containing feature importance and accuracy for the random forest model.

For datasets called with the GATK4 toolset, you can expect a similar feature importance profile. Overall ***test_accuracy*** should be about 0.99.

#### Rank and bin variants

Copy the RF model ID from the previous step into the config file, as the value for the "`rf_model_id`" key and run all steps to the end of the variant QC stage. This will generate a random forest (RF) score for every variant. Step 3.7 will generate a lot of warnings in the Bokeh library, like the following:

```
wxs-qc/3-variant_qc/7-plot_rf_output.py:553: FutureWarning: Series.__getitem__ treating keys as positions is deprecated. In a future version, integer keys will always be treated as labels (consistent with DataFrame behaviour). To access a value by position, use `ser.iloc[pos]`
```

You can ignore these warnings.

#### Variant QC results and interpretation

The RF model attributes to each variant a score value (between 0 and 1), which describes the confidence level that this variant is a true positive. However, working with individual scores is complicated. Therefore, we split the scoring interval into 100 bins and assign each variant to an RF bin. **Note:** for historical reasons, the lower RF bins correspond to more confident values. For example, bin 45 contains higher confidence variants than bin 95. The goal of this step is to choose a possible RF bin cutoff: this cutoff will be re-evaluated in stage 4, and variants with RF score greater than the chosen cutoff will be discarded in that stage.

After assigning the RF bin, we plot several variant-level metrics depending on the RF bin. All these plots follow the same structure: The X-axis is the RF bin. The Y-axis is the metric value. Each graph has two versions:

- **Simple** - Y-axis shows the raw metric value for each bin. The point size represents the number of variants in the bin.
- **Cumulative** - Y-axis shows the sum of values for the current bin and all smaller-value bins (containing more confident variants). That is, we count all variants with bins lower than the value on the X-axis and don't count all variants with higher RF bins.

We will concentrate on the cumulative plots.

To estimate the overall quality of the RF model, we have the following graphs:

- Number of SNVs and indels failing hard filters - these should be FP variants (Figure 9).
- Number of variants from known high-quality datasets - these should be TP variants (Figure 10).

In an ideal world, we'd like to have a model that classifies all TPs as high-confidence and puts them in the lowest bin (01-02), and at the same time classifies all FPs as low-confidence and puts them in the highest bin (99-100). In reality, some TPs in our dataset can be located in low-coverage regions or appear in a small subset of samples, and therefore will have a high RF bin. In addition, some real variants can be located in low-coverage areas and thus can be marked as FPs. Thus, on graphs, we'll see that with an increase in the RF bin (and therefore a more relaxed filtering criteria), the number of both FPs (failing hard filters) and TPs (variants from well-known datasets) will increase. What we want to do is to find a threshold(s) where we have the highest possible number of passed TPs and at the same time the minimum number of passed FPs. You can find a more detailed description and analysis in (Koko *et al.* 2024), section "Generating quality scores for variant QC".

For FPs, in Figure 9, you should see the elbow point after which the number of FP variants starts to rise. To know the exact bin, zoom in and hover the mouse over the bin label. For the example dataset, this **is bin 92 for SNVs.** For indels, we have two elbow points: bins 74 and 85. We can choose bin 80 as a provisional middle point. Remember these "elbow" bins. In stage 4, we'll use them as reference points to select the hard filter evaluation intervals.

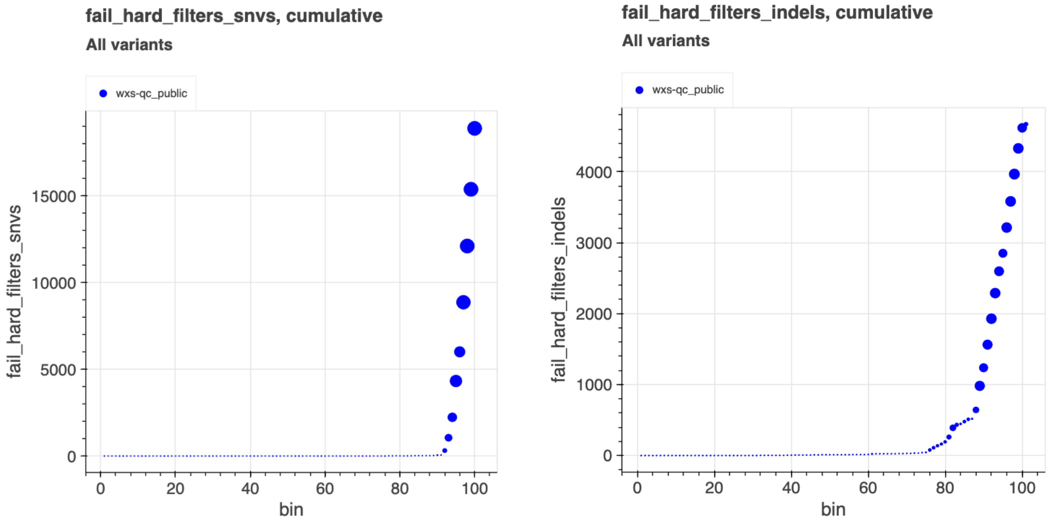

Figure 9. Number of SNVs failing at least one of the hardfilters, depending on the RF bin threshold.

For TPs (Figure 10), the number of 1000 genome SNVs reaches the maximum value also at bin 90-92. The OMNI and HapMap SNV graphs are not so clear, but the number of remaining variants on bin 92 is very close to the maximum. The number of remaining variants from the Mills and 1000G gold standard indels dataset is also close to the maximum. Therefore, we can use bin 92 as a provisional threshold point, with high number of TPs and small number of FPs.

Next, check that the Ti/Tv ratio and the ratio of transmitted to untransmitted singletons are as expected (Figure 11). Note that we can compute the transmitted / untransmitted singleton ratio because we have trios in the example dataset. If you don't have trios, this plot will not appear. For the chosen elbow point RF bin thresholds (92 for SNVs and 80 for indels) the Ti/Tv ratio is close to 2.5, and the transmitted/untransmitted ratio is about 1.

You can use the variant QC results to apply the filtering on the variant level and export data. However, the WxS-QC pipeline implements a novel hardfilter evaluation step, which allows you to analyse the combined effect of variant-level and genotype-level filtering. Therefore, remember the chosen RF cutoffs for SNVs and indels, and let's proceed to the genotyping QC stage.

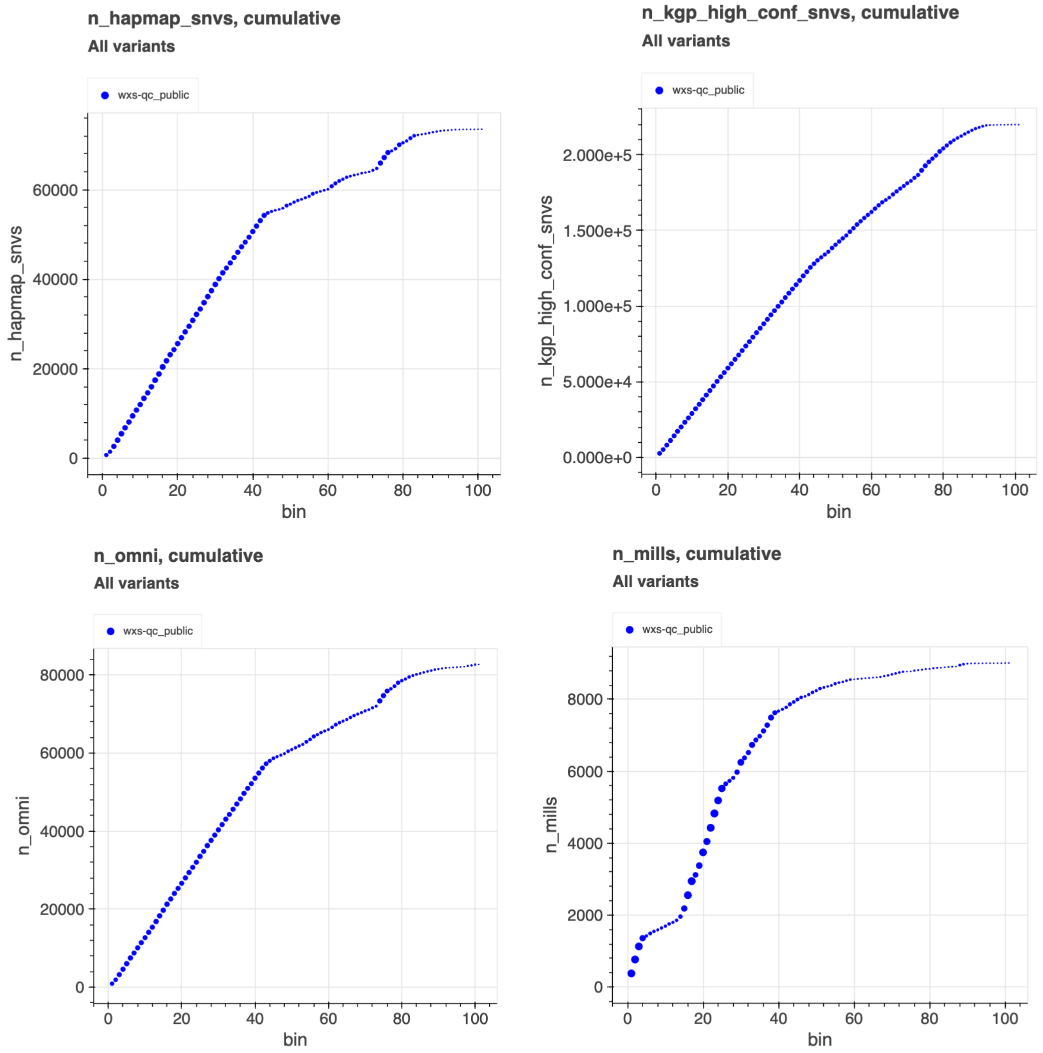


Figure 10. The number of variants from the GATK resource bundle (*GATK Resource Bundle* 2025) that remain in the example dataset, depending on the RF bin threshold. The abbreviations in the Y axis:

- `kgp_high_conf_snvs`– SNVs from 1000 Genomes (1000 Genomes Project Consortium *et al.* 2015),
- `hapmap_snvs` – SNVs from the HapMap (Frazer *et al.* 2007) project,
- `mills` - Mills and 1000G gold standard indels,
- `omni` - OMNI 2.5 genotypes for 1000 genomes samples and sites.

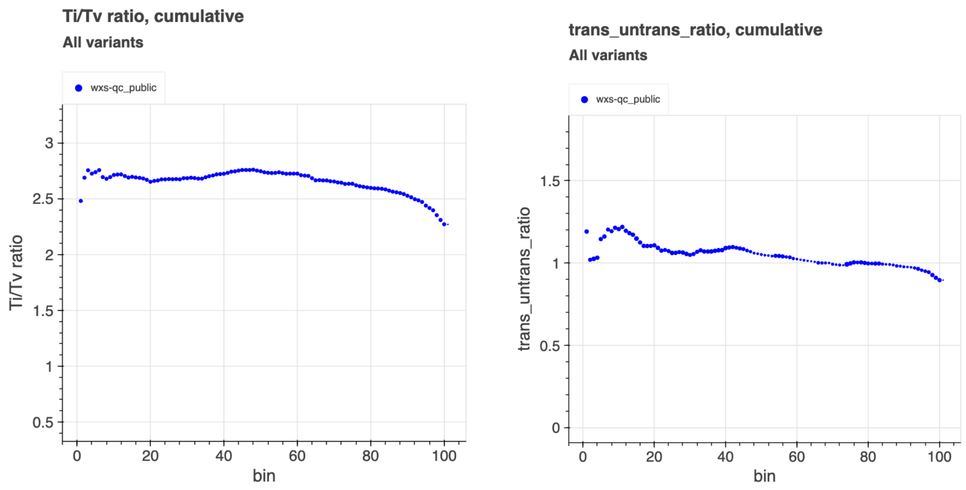


Figure 11. The Ti/Tv ratio and the ratio of transmitted to untransmitted singletons depend on the RF bin threshold.

### Stage 4. Genotype QC

The first step of the genotype QC is the hard filter evaluation. In this step, we test different combinations of hard filters and choose optimal values for each hard filter. In addition to the random forest bin that we used to rank variants, we evaluate the following genotype-level hard filters: Genotype quality (GQ), read depth (DP), and allele balance for heterozygotes (HetAB). The calculation process is described in the pipeline howto. Please refer to it for the explanation of all filters that we apply and metrics that we use. Briefly, for each hardfilter combination, we do the following:

1. Make a copy of the original dataset to work with.
2. Apply RF bin threshold and remove all variants that do not pass the threshold.
3. Apply genotype-level filters and remove all genotypes that do not pass thresholds
4. For each variant, we calculate the call rate — the fraction of samples where a genotype is called at this position.
5. Remove all variants that do not pass the call rate threshold.

By testing combinations of variant-level and genotype-level hard filters, we increase our chances of finding the best combination that keeps as many true positive variants as possible and filters out most of the false-positive ones.

However, testing combinations is the most time-consuming part of the analysis. Therefore, we recommend that you test a few combinations at the first pass, review the results and then add more combinations if needed. The calculation script caches the results, so you reuse all previously calculated combinations.

#### Evaluate hardfilter combination - round 1

In round 1, we will re-evaluate the RF cutoff chosen at the previous stage, and check its performance with additional genotype-level filtering.

At first, we recommend selecting only a few bins. If the **b** is the RF bin cutoff from the variant QC stage (see Figure 9), we use (**b**-4, **b**-2, **b**, **b**+2, **b**+4) for SNVs and (**b**-8, **b**-4, **b**, **b**+4, **b**+8) for indels.

Keep all other hardfilters at minimum default values: DP=5, GQ=10, hetAB=0.2, call_rate=0.5. For the example dataset, we run the evaluation with the extended set of bins for SNV and indels. Also, we use the config flag ***evaluate_unfiltered*** to evaluate results for unfiltered data.

Run the hardfilter evaluation and explore the resulting graphs. All evaluation graphs are interactive and allow for filtering of data points. You can switch on/off a particular hard filter and see how the graph changes. Alternatively, you can take the results data from the annotation folder and visualise it in an external tool.

The first graph to evaluate is the TP/FP plot, which shows the percentage of remaining TPs and FPs for each hard filter combination (Figure 12). For TP/FP and precision/recall, the aspect ratio is fixed to 1:1. You can disable this behaviour by a checkbox in the left part of the graph HTML page (not shown in Figure 12).

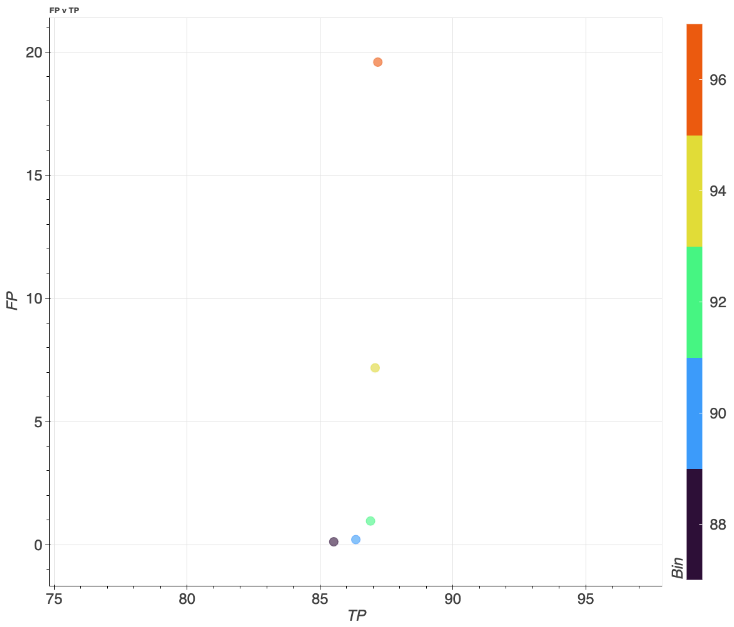


Figure 12. SNV TP/FP plot for the first round of hardfilter evaluation. All genotype-level hard filters are fixed.

The TP/FP graph shows an elbow at a similar location to the variant QC elbow: bin 92. The FP value (0.9%) is reasonable. However, for this dataset, the observed TP value (87%) is lower than that seen in (Koko *et al.* 2024). A possible explanation for the low TP value observed here is the ***call_rate*** filter. The ***call_rate*** cutoff of 0.5 is reasonable for big datasets (>1000 samples). For the example dataset, which contains only 114 samples, the call_rate cutoff is possibly too strict. We can try to achieve a better TP rate by reducing the ***call_rate*** cutoff in the next round.

#### Evaluate hardfilter combination - round 2

On the second round of calculations, we want to choose the optimum cutoffs that we will use. Therefore, we evaluate more combinations of filters:

- SNV RF bins: 86, 88, 90, 91, 92, 93, 94, 96
- Indel RF bins: 68, 72, 74, 76, 78, 80, 84, 88

- DP:  5, 10
- GQ:  10, 15
- HetAB: 0.2, 0.3
- call_rate: 0.1, 0.5

The pipeline evaluates metrics for every combination of hard filters.
Run the calculations and review the graphs after the second iteration. The generated graphs are complex because, for now, the pipeline plotting function does not support different point shapes. To analyse these data, we recommend using interactive investigation - turning on/off specific filters (DP, AB, call_rate) and observing how the data changes. You can do it using interactive capabilities of the graphs generated by the pipeline, or by opening the evaluation results tables (`hard_filter_evaluation.snv.tsv`, and `hard_filter_evaluation.indel.tsv`) in any external visualisation tool.

The interactive HTML graphs with results are available in the supplementary materials and online:

- SNV TP-FP
- SNV precision-recall
- Indel TP-FP
- Indel precision-recall

##### Precision-recall evaluation

In the second round, we consider the precision-recall graph (Figure 14) in addition to the FP/TP graph. Precision/recall is only calculated when the dataset contains a "gold-standard" sample with high-confidence intervals, where we know all real variants. For this sample, inside high-confidence intervals, we can compare our results with the expected results, calculate the number of real true-positive, false-positive and false-negative variants, and compute precision and recall.

Our example dataset contains the NA12878 (HG001) sample from the Genome in a Bottle (GIAB) project (Zook *et al.* 2016), so we can use it for precision/recall calculations. We suggest including "gold-standard" samples in any data cohort, and rely on precision/recall graphs to determine the best combination of filters.

##### SNVs

After reviewing the FP/TP (Figure 13) and the precision-recall (Figure 14) plots, we can see the following:

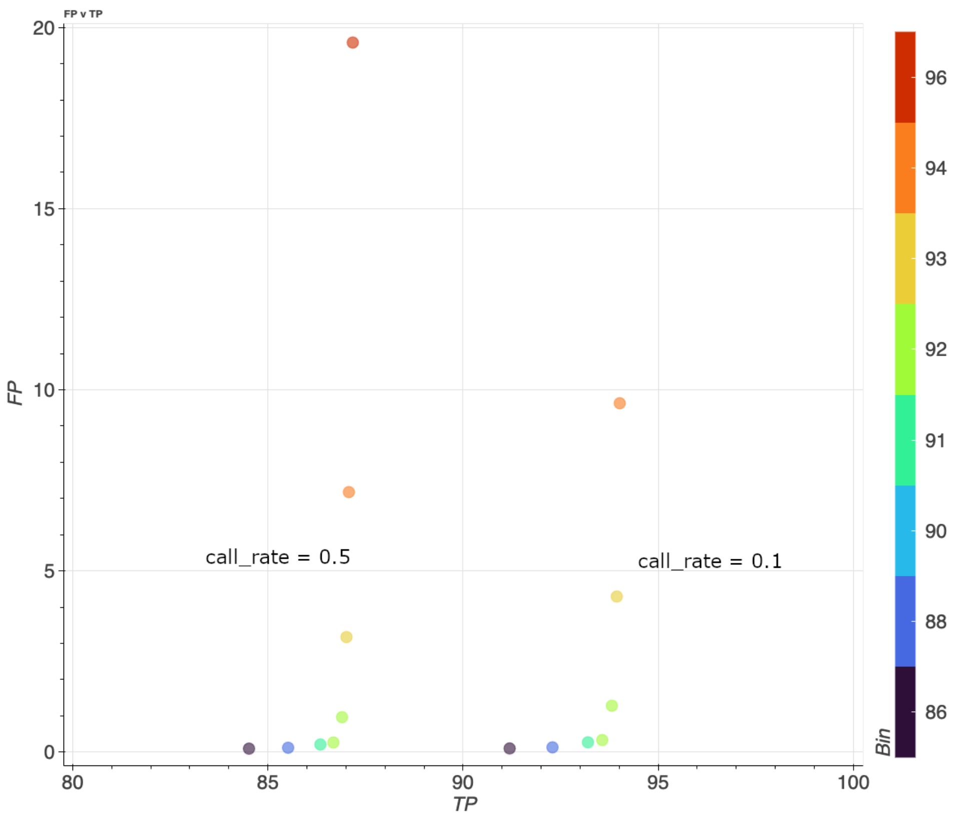


Figure 13. SNV TP/FP plot for the second round of hard filter evaluation, for two different call_rate values. All other filters are fixed: DP=5, GQ=10, HebAB=0.2

- Changing the call rate to 0.1 noticeably increases TP without a significant effect on FP, as we expected. For the precision/recall graph (Figure 14), decreasing the call rate to 0.1 increases recall without a significant impact on precision.
- In the same way, we observe that increasing the DP threshold decreases TP without a significant effect on FP (figure not included into the text, you can explore the interactive SNV TP-FP plot). Increasing the DP threshold also slightly improves precision, but decreases recall. Therefore, in this case, there is no sense in using a higher DP threshold.
- Similarly, Increasing GQ and HetAB thresholds makes sense only for RF bins >=96 (data not shown, you can explore the interactive SNV TP-FP plot)

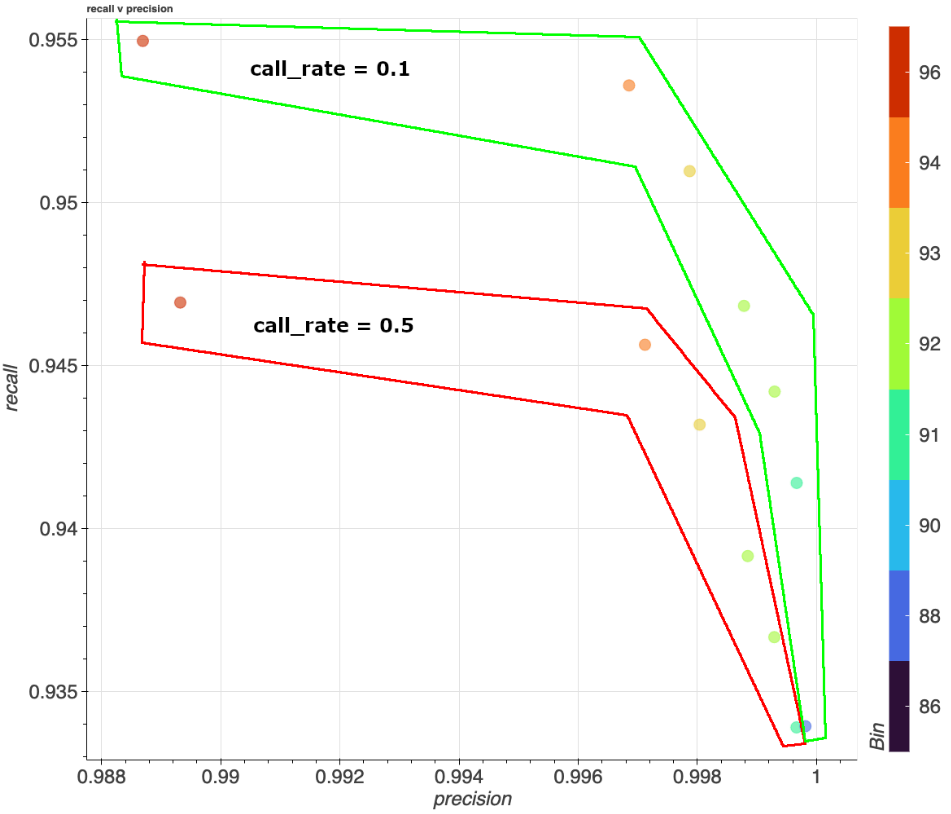


Figure 14. SNV precision/recall graph for two different call_rate values. All other filters are fixed: DP=5, GQ=10, HebAB=0.2

Check the graphs for the other metrics. For the elbow point bins (91-93) we observe the transmitted/untransmitted ratio in the range [1-1.01] (Figure 15), and the Mendelian error rate in the range $(4 - 5)*10^{-4}$ (data not shown). Increasing HetAB moves the transmitted/untransmitted ratio closer to 1 (Figure 15) – we can use it for stringent filtering.

Finally, for the example dataset, we choose three combinations of hard filters (Table 1):

- Relaxed, allowing about 10% of FP, but having as much recall as possible.
  This yields maximum recall with reasonable precision. This choice is optimal for rare variant identification.
- Medium, close to the bend point on the TP/FP graph.
- Stringent, achieving maximum precision.

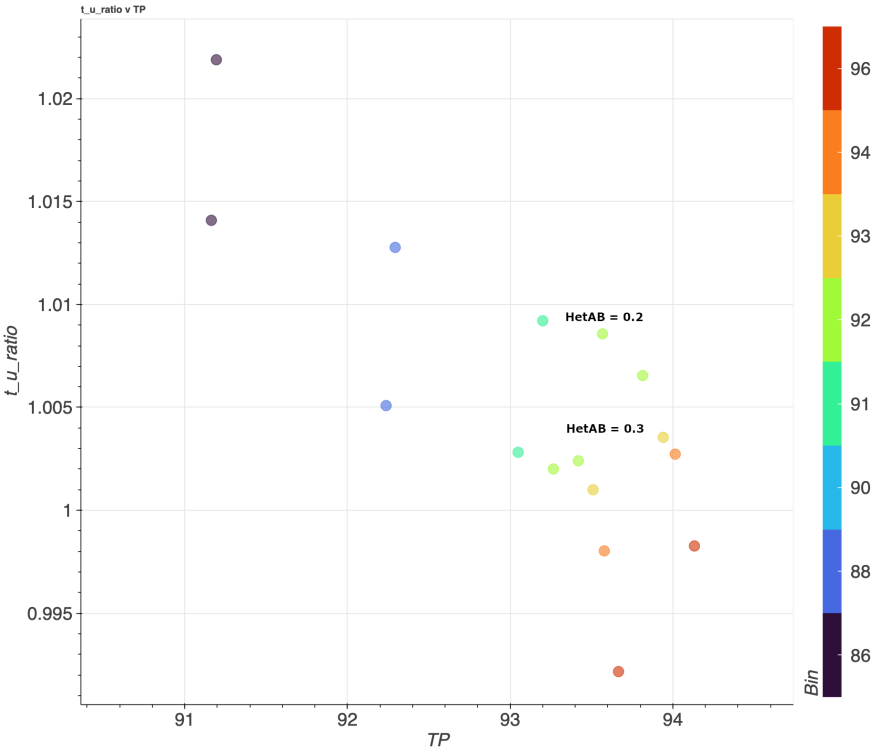


Figure 15. Transmitted/untransmitted singleton SNV ratio depending on RF bin and Het Allele Balance.

##### Indels

For indels, the best choice is more subtle. Decreasing the **call_rate** increases TP but also decreases precision (Figure 16). Increasing HetAB to 0.3 improves precision without affecting TP and with an affordable decrease in recall (data not shown), so it's a good choice for stringent filtering.

Finally, we have chosen the set of thresholds presented in Table 2.

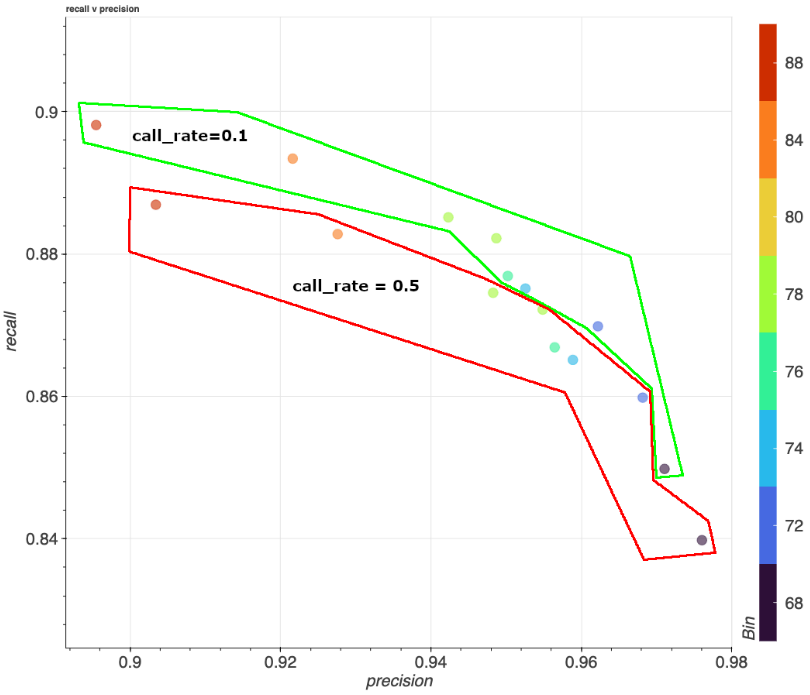


Figure 16. Precision/reacall depending on the RF bin for indels.

#### Export data to VCF format

Put the chosen hard filter combinations for the relaxed, medium and stringent filter combinations in the config file, and run genotype QC and data export (steps 4.2 and 4.3a), as described in the pipeline howto. The pipeline removes all genotypes and variations that fail the relaxed hard filter threshold, flags all variants and genotypes passing medium and stringent filters, and exports data to VCFs. For our example dataset, 9.4% of coding variants and 15.5% of non-coding variants were removed after genotype QC, because they failed the relaxed hard-filter thresholds.

Finally, you can run the mutation spectra on the final data to ensure that the mutation spectrum after QC process (Figure 17) looks reasonable.

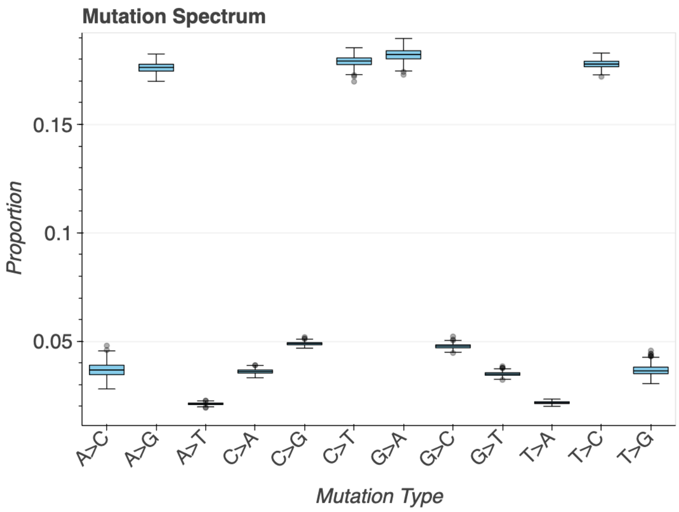


Figure 17. The example dataset mutation spectra for after-QC data.

### QC results summary and comparison

We successfully performed sample, variant and genotype QC using the WxS-QC pipeline. Now we can outline the overall QC results.

#### Sample QC

The contamination check via VerifyBamID revealed no suspicious samples.

The sex imputation and comparison (Figure 3) revealed potentially mislabeled samples. However, the mislabelled samples do not affect the subsequent sample and variant QC, therefore, we decided to label and keep them in the dataset.

Superpopulation identification and stratified filtering allow us to filter out outlier samples. This step is essential for subsequent variant QC because variants in problematic samples affect variant frequencies and metrics and can alter the results of the Variant QC.

Whereas filtering based on mean dispersion has a risk of removing good samples, for GWAS and similar types of cohort analysis, this approach produces good results. We applied it to several data cohorts that were sequenced and analysed at the Wellcome Sanger Institute (Koko *et al.* 2024).

#### The variant and genotype QC

To estimate the effect of variant and genotype quality filtering, we can compare the results of the WxS QC pipeline with those of standard genotype-based filtering using hard thresholds on the genotype metrics DP and GQ only. To implement standard filtering, we ran an evaluation without any filtering by RF bin and HetAB, and filtered out all genotypes with DP < 5 or GQ < 10 for "relaxed" filtering and DP < 10 or GQ < 20 for "stringent".

Since this standard filtering works only on the genotype level and doesn't analyse variant-level quality metrics (QD, MQ, FS), it can't effectively distinguish TPs and FPs. Therefore, we expect that many of the FPs will remain in the dataset, and the FP values for standard filtering will be high. To make the results more comparable, we also tested a combination of genotype-level and variant-level filtering with hard thresholds. For "relaxed" standard filtering, we used a single threshold variant QUALITY>=100 in addition to the genotype filters. For "stringent" standard filtering, we applied additional variant-level QD, FS and MQ thresholds.

The resulting summary table of the QC metrics for SNVs is presented in Table 1. The WxS QC "relaxed" filter achieves a precision of > 99% and a recall value of 95.36%. By comparison, the "relaxed" standard filtering achieves a precision and recall of approximately 95% and "stringent" standard filtering reduces recall to 93.9%, with a slight increase in precision (95.8%). This shows that WxS QC improves precision and recall across a range of filter criteria, in comparison to a standard hard-threshold filtering.

The combination of genotype-level standard filters and variant-level hard thresholds performs better than the genotype-only filtering. Filtering by quality >=100 decreases the number of FPs to 76%, but has little effect on precision and recall. Stringent standard filtering with QD, MQ and FS thresholds works noticeably better: It eliminates FPs completely (which is expected because we initially label variants that fail these thresholds as FPs), and achieves a precision 99.13% and a recall 93.62%. However, the WxS-QC “medium” filtering achieves better precision, recall, and TP values. The WxS QC “stringent” filtering achieves precision >=99.9% with slightly smaller recall and TP number.

Table 1. The comparison between the standard data QC approach using only DP and GQ filtering, and the WxS-QC pipeline for SNVs. The high FP values for standard filtering without variant-level filters are expected (see the explanation in the text).

| | **Standard** | | **Standard + filters** | | **WxS QC** | | |
|---|---|---|---|---|---|---|---|
| | **relaxed** | **stringent** | **relaxed** | **stringent** | **relaxed** | **medium** | **stringent** |
| Filters | - | - | QUAL >=100 | QUAL>=100 QD>=2 FS<=60 MQ>=30 | - | - | - |
| RF bin | - | - | - | - | 94 | 91 | 88 |
| DP | 5 | 10 | 5 | 10 | 5 | 5 | 5 |
| GQ | 10 | 20 | 10 | 20 | 10 | 10 | 10 |
| HetAB | - | - | - | - | 0.2 | 0.2 | 0.2 |
| Call rate | - | - | - | - | 0.1 | 0.1 | 0.1 |
| TP, % | 96.74 | 93.46 | 95.81 | 92.81 | 94.01 | 93.57 | 92.29 |
| FP, % | 97.87 | 95.66 | 76.43 | 0.00 | 9.63 | 0.33 | 0.13 |
| Mendelian error $*10^{-4}$ | 9.66 | 6.9 | 9.65 | 4.58 | 5.82 | 4.36 | 3.07 |
| Precision % | 95.53 | 95.64 | 95.65 | 99.13 | 99.68 | 99.93 | 99.98 |
| Recall, % | 95.81 | 93.86 | 95.80 | 93.62 | 95.36 | 94.42 | 93.40 |
| t/u ratio | 0.95 | 0.96 | 0.96 | 1.00 | 1.00 | 1.01 | 0.93 |

The advantages of the WxS-QC pipeline are clearer for indels (Table 2). “Relaxed” standard filtering achieves slightly better recall than the “relaxed” WxS-QC (91.05% and 89.34%), but with a significant drop in precision (73.6% vs 92.1%). The “stringent” standard filtering with variant-level thresholds achieves a precision of 80.7%, but it is still notably worse than the results of the WxS-QC pipeline (from 92 to 98% depending on the filtering settings). Similar to SNVs, the “medium” WxS-QC filtering achieves better precision, recall, and TP values, having slightly more FP variants.

Table 2. Comparison between the standard data QC approach using only DP and GQ filtering and the WxS-QC pipeline for Indels. Regarding high FP values, see the note to Table 1.

| | **Standard** | | **Standard + filters** | | **WxS QS** | | |
|---|---|---|---|---|---|---|---|
| | **relaxed** | **stringent** | **relaxed** | **stringent** | **relaxed** | **medium** | **stringent** |
| Filters | - | - | QUAL >=100 | QUAL>=100 QD>=2 FS<=60 MQ>=30 | - | - | - |
| RF bin | - | - | - | - | 84 | 78 | 72 |
| DP | 5 | 10 | - | - | 5 | 5 | 5 |
| GQ | 10 | 20 | 5 | 10 | 10 | 10 | 10 |
| HetAB | - | - | 10 | 20 | 0.2 | 0.2 | 0.3 |
| Call rate | - | - | - | - | 0.1 | 0.1 | 0.1 |
| TP, % | 94.24 | 89.84 | 93.25 | 88.59 | 89.32 | 88.63 | 87.40 |
| FP, % | 97.92 | 95.76 | 82.74 | 0 | 8.20 | 2.31 | 0.62 |
| Mendelian error $*10^{-4}$ | 149.6 | 107.31 | 154.19 | 88.93 | 78.05 | 73.49 | 55.65 |
| Precision % | 73.62 | 75.02 | 73.76 | 80.70 | 92.16 | 94.87 | 98.23 |
| Recall, % | 91.05 | 80.88 | 91.04 | 87.92 | 89.34 | 88.22 | 84.92 |

The main reason the WxS-QC pipeline achieves better results is that it filters variants using a random forest model. Filtering by the carefully selected RF bin can eliminate most of the FP variants, whilst keeping almost all TP variants. A similar approach is implemented by the GATK Variant Quality Score Recalibration (VQSR). However, implementing variant and genotype QC in the WxS-QC pipeline has several significant advantages.

- We do not rely on a "black box" solution inside a variant caller. Instead, we can choose the threshold depending on our task - to make more reliable data for common variant identification, or preserve as many variants as possible for rare variant analysis.
- The random forest model allows flexibility in the features added. For example, a part of the ALSPAC dataset sequenced at the Wellcome Sanger Institute (Koko *et al.* 2024) had a significant number of samples containing a C->A sequencing artefact. By marking all C->A SNVs with a boolean flag and adding the flag to the random forest features, we eliminated most of the artefact variations without an expensive resequencing of the dataset.
- The hard filter evaluation step in Genotype QC, especially when precision/recall calculations are enabled, provides us with even more control over this process. We can observe the percentage of filtered variants, analyse different metrics, and use genotype-level filters to improve the results compared to the standard genotype-only filtering.

In summary, the WxS-QC pipeline can achieve higher precision values while maintaining similar recall values to standard genotype and variant filtering approaches. This is especially true for indels, where the WxS-QC pipeline achieves notably higher precision values. Using standard genotype filtering could be appropriate for high-sensitivity experiments, where we aim to call as many variants as possible, even at the cost of higher false-positive rates.

#### Conclusion

The WxS-QC pipeline is a powerful genetic variant QC solution that provides users with flexibility and precise control over the genome/exome data QC process. The implemented pipeline achieves higher precision levels than standard threshold-based filtering with a comparable recall level, especially for indels.

### The WxS-QC example dataset

The public example dataset we use to test the WxS-QC pipeline is prepared from a subset of open 1000 genomes samples, and is freely available for download from https://wxs-qc-data.cog.sanger.ac.uk/wxs-qc_public_dataset_v3.tar. The dataset is designed to cover all super populations and mimic outliers in real experimental datasets.

This dataset contains the combined VCF file `joint_germline.vcf.gz` and the set of metadata files:

- `verify_bam_id_result.selfSM.tsv` - VerifyBamID results
- `self_reported_sex.tsv` - self-reported sex for all 1000 genomes samples.
- `trios.fam`, `trios-withparents.fam` – pedigree information. The pipeline now uses the version without parents. Switching to the full version with parents is in progress.
- `all_consequences_with_gene_and_csq.tsv.bgz`, `all_consequences.tsv.bgz`, `synonymous_variants.tsv.bgz` - VEP annotation in different formats. For now, the pipeline uses it all. The work on unifying the VEP input is in progress.

In the `dataset-build` subfolder, you can find all the scripts we used to prepare the data. The scripts are numbered and contain all the details to trace data preparation processes and ensure reproducibility.

Briefly, the preparation process is the following:

1. Download the pre-selected set of high-coverage CRAMS from 1000 genomes FTP (https://ftp.1000genomes.ebi.ac.uk/vol1/ftp/data_collections/1000_genomes_project/data/, released 2015-12-22).
2. Run Nf-Core Sarek pipeline (https://github.com/nf-core/sarek) with GATK4 HaplotypeCaller to call per-sample variations and make joint-calling. The resulting VCF is the initial data for the QC pipeline.
3. Run VerifyBamID on CRAMs and collect FreeMix scores.
4. Download the file with self-reported sex information and change the header to match the pipeline requirements.

5. Run VEP on the VCF to obtain consequences. To speed up calculations, we normalize the VCF and split it into several chunks.


Temporary table to compare results
